# Evolutionary coupling range varies widely among enzymes depending on selection pressure

**DOI:** 10.1101/2020.12.19.423588

**Authors:** Julian Echave

## Abstract

Recent studies proposed that enzyme active sites induce evolutionary constraints at long distances. The physical origin of such long-range evolutionary coupling is unknown. Here, I use a recent biophysical model of evolution to study the relationship between physical and evolutionary couplings on a diverse data set of monomeric enzymes. I show that evolutionary coupling is not universally long-range. Rather, range varies widely among enzymes, from 2Å to 20Å. Furthermore, the evolutionary coupling range of an enzyme does not inform on the underlying physical coupling, which is short-range for all enzymes. Rather, evolutionary coupling range is determined by functional selection pressure.

**SIGNIFICANCE:** Until recently, only residues near enzyme active sites were thought to be evolutionarily constrained. However, recent studies proposed that active sites induce long-range evolutionary constraints. This seems to conflict with the common finding that physical couplings in proteins are short-range. This raises the question of how short-range physical couplings may cause long-range evolutionary couplings. Here, I show that the function that maps physical coupling into evolutionary coupling depends on functional selection pressure. Under weak selection, both couplings are similarly short-range; under strong selection, short-range physical coupling is non-linearly turned into long-range evolutionary coupling. Thus, due to a huge variation of selection pressure, evolutionary coupling range varies widely among enzymes, from very short (2 Å) to very long (20 Å).

---

As enzymes evolve, different sites evolve at different rates. The main reason for such variation of evolutionary rate among sites within proteins is selection for stability (1–4). Until recently, activity constraints were thought to affect just the few residues directly involved in catalysis and their immediate neighbours (1, 5). However, recent studies have reported that active sites influence evolutionary rates at long distances, slowing down the evolution of residues as distant as 30 Å (6–10). It seems reasonable to assume that such long-range evolutionary coupling could result from long-range physical couplings, such as those involved in allosteric mutations (11, 12). This would align with the notion that enzymes are evolutionarily designed to optimize long-range coupling (13–15). However, except perhaps for a few allosteric residues, I would expect physical coupling to be a be the typical short-range exponentially decreasing function of distance characteristic of indirect through-the-contact-network couplings (14, 16–20). If this is the case, it would leave long-range evolutionary coupling begging explanation.

The aim of this work is to verify whether physical coupling is short-range and, in that case, to study how such a short-range physical coupling may give rise to a long-range evolutionary coupling. To this end, I use the stability-activity model of enzyme evolution, MSA (21). I previously showed that this model reproduces quantitatively the observed slow increase of evolutionary rate with distance from the active site that led to the proposal of long-range evolutionary couplings (7). This makes MSA suitable for exploring the physical underpinnings of long-range evolutionary coupling.

Before describing the model, I start with some definitions. The evolutionary rate, *K*, is the number of amino acid substitutions per unit time along an evolutionary trajectory; *K*_0_ is the rate for the case in which all mutations are neutral; 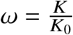 is the rate relative to the neutral rate; and 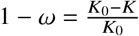 is the relative slowdown with respect to the neutral evolution case. As selection pressure increases, evolution slows down: *K* and *ω* decrease, and 1 − *ω*, the relative slowdown, increases. *K*, *ω*, and 1 − *ω* contain exactly the same information. Therefore, in what follows, to measure the effect of selection on evolution, I will mostly use 1 − *ω*, the evolutionary slowdown.

With the help of the MSA model, I define physical and evolutionary coupling measures, and derive the formula that relates them. (The MSA model is described in detail in the Supporting Material document (SM), in SM Section 1 and SM Section 3, and in (21).) MSA predicts that the evolutionary slowdown of an enzyme residue *r* due to selection on stability and activity is given by (Eq. S19):

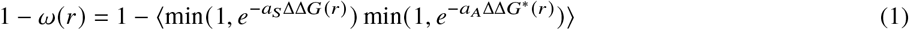

where ΔΔ*G*(*r*) and ΔΔ*G*\*(*r*) are, respectively, mutational changes of folding free energy and activation free energy; *a*_*S*_ and *a*_*A*_ are positive parameters that represent selection pressure on stability and activity, respectively; and ⟨⋯⟩ stands for averaging over mutations. Eq. 1 relates the evolutionary slowdown of residue *r* to the effects of mutating this residue on stability and activity. Since here I am only interested on selection on activity, I consider a hypothetical scenario in which selection on stability is turned off. Replacing *a*_*S*_ = 0 into Eq. 1 it follows that the evolutionary slowdown of site *r* due to selection on activity is given by (Eq. S20):

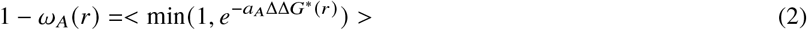

The mutational activation free-energy change ΔΔ*G*\*(*r*) is due to the distortion of the active-site caused by mutating residue *r* (21) (Eq. S45, SM Section 1.1, and SM Section 3.3). Therefore, ΔΔ*G*\*(*r*) represents the *physical coupling* between the enzyme’s active site and residue *r*. This causes the slowdown 1 − *ω*_*A*_(*r*), which, therefore, can be considered a measure of *evolutionary coupling*. Thus, Eq. 2 governs how evolutionary coupling (1 − *ω*_*A*_) depends on physical coupling (ΔΔ*G*\*) and functional selection pressure (*a*_*A*_). (For notational simplicity, I will drop the explicit reference to residue *r* whenever possible.)

I studied the relationship between physical and evolutionary couplings on a data set of monomeric enzymes used previously (21) (SM Section 1.2). Briefly, for each protein, I calculated ΔΔ*G* and ΔΔ*G*\* for all mutations at all residues using the Linearly Forced Elastic Network Model (LFENM). Then, I obtained the model parameters *a*_*S*_ and *a*_*A*_ by fitting MSA predictions to empirical rates. Finally, I calculated the residue-dependent evolutionary couplings, 1 − *ω*_*A*_, and physical couplings, ⟨ΔΔ*G*\*⟩. (⟨ΔΔ*G*\*(*r*)⟩ is the average of ΔΔ*G*\*(*r*) over mutations at *r*.) For details of the calculation, see SM Section 1.1. I consider MSA to be validated in a previous study (21). However, for completeness, in SM Section 2 I show the excellent agreement between MSA predictions and empirical rates (SM Section 2.1) and I discuss the adequacy of using LFENM to calculate ΔΔ*G* and ΔΔ*G*\* (SM Section 2.2). In what follows, I focus on the study of physical and evolutionary couplings.

For clarity, I start by considering three illustrative examples (Figure 1). I measure coupling range using *d*_1/2_, the distance at which coupling is half the maximum. Physical coupling (⟨ΔΔ*G*\*⟩) is similar for the three cases, it decreases exponentially with increasing distance, and it is very short range (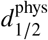 is 2.1 Å, 2.0 Å, and 2.0 Å for 1OYG, 1QK2, and 1PMI, respectively, Figure 1A). In contrast, evolutionary coupling (1 − *ω*_*A*_) varies among the examples, it is not an exponential but a sigmoid, and its range varies widely (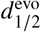 is 5.4 Å, 8.7 Å, and 15.6 Å for 1OYG, 1QK2, and 1PMI, respectively; Figure 1B). It will shown below that this variation of evolutionary coupling range is due to the variation of functional selection pressure acting on these three enzymes (*a*_*A*_, is 29.2, 80.2, and 800 for 1OYG, 1QK2, and 1PMI, respectively).

**Figure 1:**
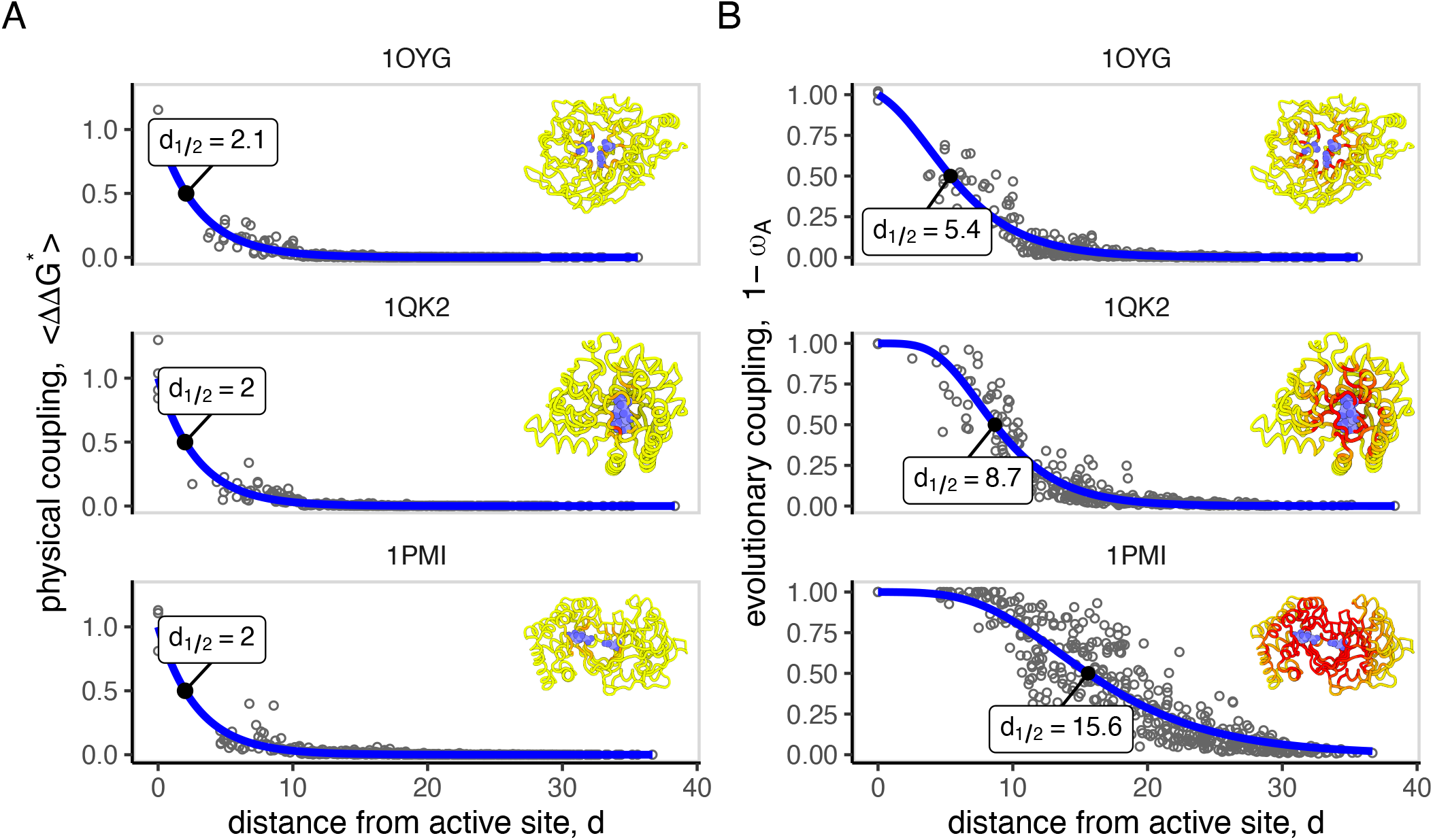
Coupling between the active site and other residues for three illustrative examples. The three cases shown are the enzymes with PDB IDs 1OYG, 1QK2, and 1PMI. **A**: physical coupling, as measured by the change in activation free energy that results from mutations; each point corresponds to a protein site; a site’s ⟨ΔΔ*G*\*⟩ is the average of ΔΔ*G*\* over mutations; the smooth line is an exponential fit 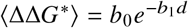, where *d* is the distance from the closest active-site residue, measured in Å; the point for which coupling is 1/2 of its maximum is displayed in black; the insets show the 3D protein structures coloured from yellow to red according to increasing physical coupling. **B**: evolutionary coupling, as measured by 1 − *ω*_*A*_, the relative slowdown of evolution due to selection on activity; each point corresponds to a protein site; the smooth line is a function 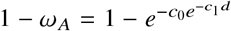, fit to the points; the point for which coupling is 1/2 of the maximum is displayed in black; the insets show the 3D protein structures coloured from yellow to red according to increasing evolutionary coupling. For the sake of comparison, couplings are scaled so that the smooth fits are 1 at *d* = 0. 3D images were made with https://3dproteinimaging.com/protein-imager (38).

Figure 2 shows the distance-dependence of physical and evolutionary couplings for all the enzymes studied. Physical coupling is a short-range exponential decline with distance, very similar for all cases (Figure 2A). In contrast, the distance-dependence of evolutionary coupling varies among proteins, from a short range exponential decline to a long-range sigmoidal decline (Figure 2B). Because physical coupling is very similar for all enzymes, from Eq. 2 it follows that the variation among enzymes of evolutionary coupling must be determined by the selection parameter *a*_*A*_. This is confirmed in F igure 2C, that shows that evolutionary coupling range is independent of physical coupling range, and Figure 2D, that shows that the variation of evolutionary coupling range is almost completely explained by the selection pressure parameter *a*_*A*_.

**Figure 2:**
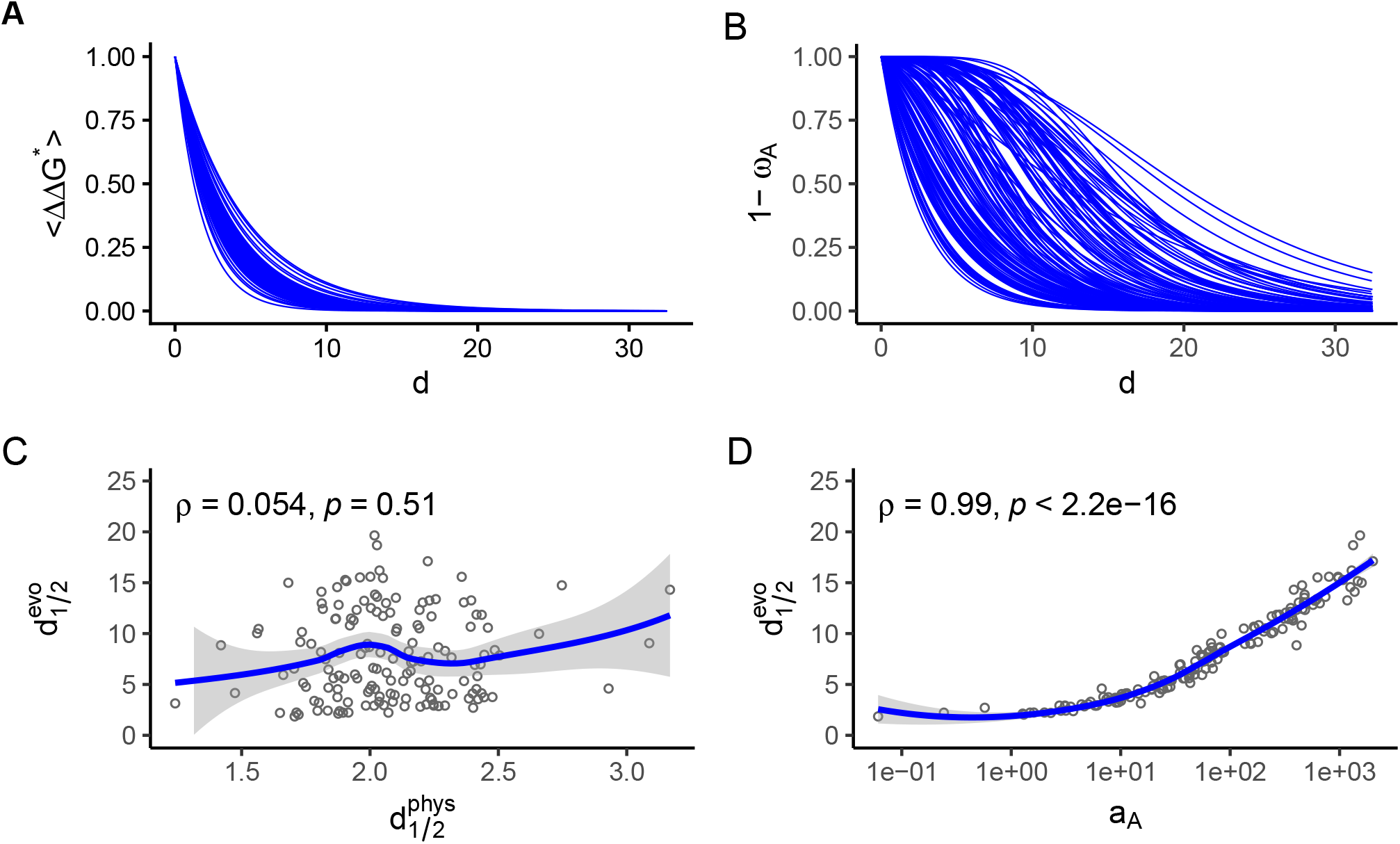
Evolutionary coupling range increases with selection pressure on activity. Coupling between the active site and other residues for each protein of a data set of 157 monomeric enzymes of diverse sizes, structures, and functions. **A**: physical coupling, as measured by ⟨ΔΔ*G*\*⟩, the activation free energy change averaged over mutations at the mutated site; each line is the smooth fit 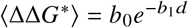 for one protein of the data set, where *d* is the distance from the closest active-site residue in Å. **B**: evolutionary coupling, as measured by 1 − *ω*_*A*_, the relative slowdown due to selection on activity; each line is the smooth fit 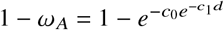 for one protein of the data set; Couplings are scaled so that all smooth fits are 1 at *d* = 0. **C**: range of evolutionary coupling, measured by 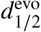, the distance at which 1 − *ω*_*A*_ becomes half of its maximum, vs. range of physical coupling, measured by 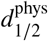, the distance at which ⟨ΔΔ*G*\*⟩ becomes half of its maximum. **D**: range of evolutionary coupling, measured by 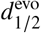, the distance at which *d*, vs. functional selection pressure, measured by model parameter *a*_*A*_. In **C** and **D**, each point represents one protein, *ρ* is Spearman’s correlation coefficient, *p* its p-value, and the lines are local regression fits.

Thus, according to Figure 2, evolutionary coupling range varies widely among enzymes, from 1.9 Å to 19.7 Å, as a result of the variation of parameter *a*_*A*_ over more than 4 orders of magnitude, from 6 × 10−2 to 2 × 10^3^. This huge variation of *a*_*A*_ represents a variation of the functional selection pressure under which enzymes evolve (SM Section 2.3). This variation can be explained by Eq. 2, that non-linearly maps a short-range exponentially decreasing physical coupling into a sigmoidally decreasing evolutionary coupling whose range increases with selection pressure (SM Section 2.4). In summary, the fundamental finding of this work is that long-range evolutionary coupling is not due to enzymes being particularly well-designed for long-range physical coupling, but it is a consequence of a non-linear amplification of physical coupling under strong functional selection pressure.

The previous findings provide a mechanism that explains how functional constraints slow down enzyme evolution. *A priori*, the decrease of the rate of evolution with increasing selection pressure could be uniformly distributed among all residues. However, the present findings imply a different picture: increasing functional selection pressure increases the range of influence of the enzyme’s active site on other residues (Figure 2 D); as this range increases, more sites become functionally constrained (Figure 3A) and the active site becomes more tightly coupled to the rest of the protein (Figure 3 B); as a result, the enzyme’s evolution slows down (Figure 3 C). In this work, I have derived *a*_*A*_ from patterns of rate variation among sites within proteins. However, the model predicts that *a*_*A*_ should also influence rate variation among proteins, which would connect rate variation within proteins with rate variation among proteins. Further work is needed to test this important prediction.

**Figure 3:**
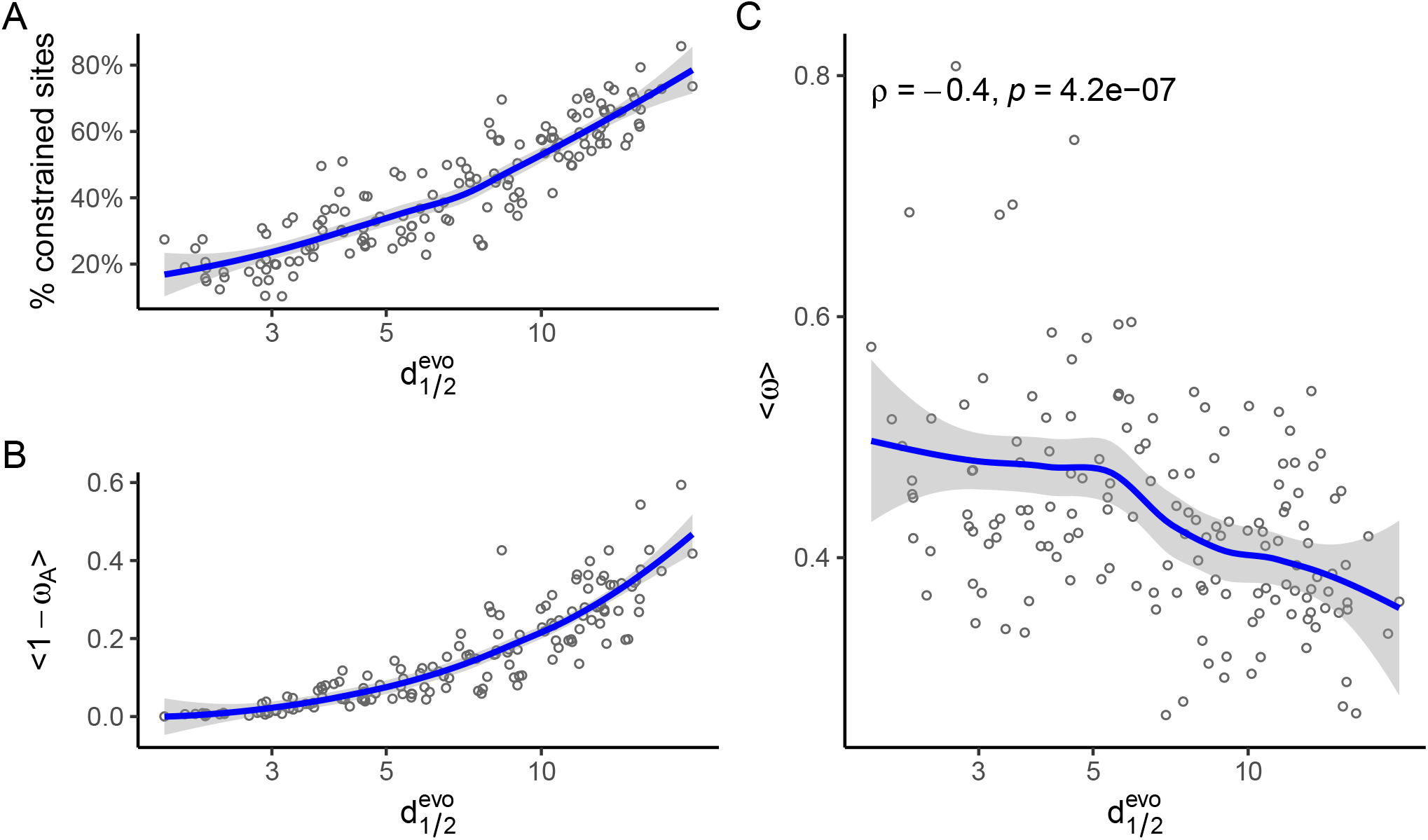
Increasing evolutionary coupling range slows down enzyme evolution. Evolutionary coupling range is measured by 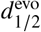, the distance at which evolutionary coupling is half its maximum. **A**: Increase of the fraction of functionally constrained sites; the number of activity-constrained residues was calculated using 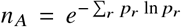, where *r* denotes residue and 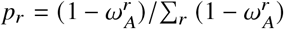. So defined, *n*_*A*_, which varies between 1 and the total number of sites, measures how distributed over sites 1 − *ω*_*A*_ is. **B**: Increase of overall enzyme coupling, measured by 1 − *ω*_*A*_ averaged over residues. **C**: Predicted decrease of the protein rate of evolution, measured by the relative rate *ω* averaged over residues; *ρ* is Spearman’s correlation coefficient, *p* its p-value. In all panels, each point represents one protein and lines are local regression fits.

To finish, I mention another two research directions suggested by this work. First, for enzymes, functional selection pressure depends on metabolic role (22–25). Specifically, the main functional constraints are enzyme-specific metabolic flow and enzyme essentiality (23). Therefore, these properties should correlate with parameter *a*_*A*_ of the present work, and, as a consequence, metabolic role should affect evolutionary coupling range. This prediction should be verified. Second, the present findings indicate the intriguing possibility of manipulating evolutionary coupling range by adjusting selection pressure, which could be explored using enzyme evolution experiments.

## SUPPORTING CITATIONS

References (26–37)

## SUPPORTING MATERIAL

See supporting_material.pdf.

## ACKNOWLEDGEMENTS

This work was supported by Consejo Nacional de Investigaciones Científicas y Técnicas (grant number PIP 112 201501 00385 CO) and by Agencia Nacional de Promoción Científica y Tecnológica (grant number PICT-2016-4209).

## DATA AVAILABILITY

The data and code underlying this article are available in Zenodo, at http://doi.org/10.5281/zenodo.4309233

