## Supplementary Information for "Evolutionary coupling range varies widely among enzymes depending on selection pressure"

Julian Echave

Instituto de Ciencias Físicas, UNSAM and CONICET, San Martín, Buenos Aires, Argentina.

### 1 Supplementary Methods

#### 1.1 Summary of the MSA model

This section summarises the stability-activity enzyme evolution model, MSA, and gathers its key equations. A more detailed description of the model and derivation of the equations is presented in Section 3 and in [1]. To avoid duplication, I use the equation numbers of Section 3, where the formulas are derived.

**Rates and slowdowns, definitions** The evolutionary rate, $K$, is the number of amino-acid substitutions per unit time along an evolutionary trajectory; $K_0$ is the rate for the case in which all mutations are neutral; $\omega = \frac{K}{K_0}$ is the rate relative to the neutral rate; and $1 - \omega = \frac{K_0 - K}{K_0}$ is the relative slowdown with respect to the neutral evolution case. As selection pressure increases, evolution slows down: $K$ and $\omega$ decrease, while $1 - \omega$ increases. To measure the effect of selection on evolution, I use $1 - \omega$, the evolutionary slowdown.

**MSA slowdowns** MSA predicts that the relative slowdown of residue $r$ is given by:

$$1 - \omega(r) = 1 - \langle \min(1, e^{-a_S \Delta\Delta G(r)}) \min(1, e^{-a_A \Delta\Delta G^*(r)}) \rangle \tag{S19}$$

where $\Delta\Delta G(r)$ and $\Delta\Delta G^*(r)$ are, respectively, mutational changes of folding free energy and activation free energy; $a_S$ and $a_A$ are site-independent positive parameters that represent selection pressure on stability and activity, respectively; and $\langle \cdots \rangle$ stands for averaging over mutations at site $r$. Eq. S19 relates the slowdown of the evolution of a site to the effects of mutating this site on stability and activity.

**Physical and evolutionary couplings** In a hypothetical scenario in which selection on stability is turned off, the slowdown caused by selection on activity is given by:

$$1 - \omega_A(r) = 1 - \langle \min(1, e^{-a_A \Delta\Delta G^*(r)}) \rangle \tag{S20}$$

where $\Delta\Delta G^*(r)$ represents the *physical coupling* between residue $r$ and the enzyme's active site; $1 - \omega_A(r)$ represents the *evolutionary coupling* between residue $r$ and the active site; Eq. S20 relates these two couplings.

**LFENM calculation of $\Delta\Delta G$** To calculate $\Delta\Delta G$ and $\Delta\Delta G^*$, I used the Linearly Forced Elastic Network Model (LFENM) [2, 3]. LFENM represents a given protein as an elastic network of nodes (amino acids) connected by harmonic springs (interactions) and models mutations as random perturbations of the lengths of the springs that connect the mutated site to other sites. The elastic network has a quadratic energy function with a single minimum at the equilibrium conformation. (A protein conformation is represented by $\mathbf{r}$, the vector of Cartesian coordinates of all network nodes.) Given a wild type protein with energy $V_{\text{wt}}(\mathbf{r})$ with minimum at $\mathbf{r}^0_{\text{wt}}$ and a mutant with energy $V_{\text{mut}}(\mathbf{r})$ with minimum at $\mathbf{r}^0_{\text{mut}}$. The difference between the equilibrium conformations is given by:

$$\mathbf{r}^0_{\text{mut}} - \mathbf{r}^0_{\text{wt}} = -\mathbf{K}^{-1}\mathbf{f} \tag{S25}$$

where $\mathbf{f}$ is a "force" vector related to the perturbations applied to model the mutation. The free-energy change caused by the mutation is:

$$\Delta\Delta G_{\gamma\gamma_0}(r) = \frac{1}{2}\sum_{j<i} k_{ij}\delta l_{ij}^2 - \frac{1}{2}(\mathbf{r}^0_{\text{mut}} - \mathbf{r}^0_{\text{wt}})^T \mathbf{K} (\mathbf{r}^0_{\text{mut}} - \mathbf{r}^0_{\text{wt}}) \tag{S28}$$

where $k_{ij}$ is the force constant of the spring connecting sites $i$ and $j$, $\delta l_{ij}$ is the change of the length of spring $ij$ due to the mutation from wild-type state $\gamma_0$ to the mutated state $\gamma$, and $\mathbf{K}$ is the Hessian matrix common to $V_{\text{mut}}(\mathbf{r})$ and $V_{\text{wt}}(\mathbf{r})$. $\Delta\Delta G$ given by Eq. S28 is the difference between the mutant's and wild-type's minimum energies (see Figure S8). Using these, Eq. S29 allows the calculation for arbitrary $\alpha \to \beta$ mutations.

**LFENM calculation of $\Delta\Delta G^*$** To calculate $\Delta\Delta G^*$ I made the following considerations [1]. First, at low substrate concentrations, the activation energy barrier is the free energy difference between the transition state $ES^*$ and enzyme $E$ and substrate $S$ free in solution: $\Delta G^* = G(ES^*) - G(E) - G(S)$. Second, $\Delta G^*$ can be written as the sum of two contributions, a *distortion energy*, which is the energy necessary for the enzyme and substrate to adopt their transition state conformations, $E^*$ and $S^*$, plus a *vertical binding energy*, which is the energy released when $E^*$ and $S^*$ bind to form $ES^*$. Third, I assume that mutations have no effect on the vertical binding energy or the substrate distortion, but affect only the enzyme's distortion free energy: $\Delta\Delta G^* = \Delta G(E \to E^*)$. Finally, I assume that the wild-type has a preorganized active site with a conformation which is identical to the transition state conformation. Using these assumptions and the LFENM model, I derived that $\Delta\Delta G^*$ is the energy needed to distort the mutant's active site from the mutant's conformation $\mathbf{r}^0_{a,\text{mut}}$ to the wild type conformation $\mathbf{r}^0_{a,\text{wt}}$ (Figure S8), which is given by:

$$\Delta\Delta G^*_{\gamma\gamma_0}(r) = \frac{1}{2}(\mathbf{r}^0_{a,\text{wt}} - \mathbf{r}^0_{a,\text{mut}})^T \mathbf{K}^{\text{eff}}_{aa} (\mathbf{r}^0_{a,\text{wt}} - \mathbf{r}^0_{a,\text{mut}}) \tag{S45}$$

where $\mathbf{K}^{\text{eff}}_{aa}$ is a matrix that allows the calculation of the effective energy of distortions within the conformational subspace spanned by active residue coordinates $\mathbf{r}_a$. Eq. S45 corresponds to mutations starting at wild-type state $\gamma_0$; From these, Eq. S46 allows the calculation of other $\alpha \to \beta$ mutations.

#### 1.2 Model parameters

**Elastic Network Model** There is a large variety of ENMs. Here, I used the ENM of Ming and Wall [4]: amino acids are represented by single nodes; nodes are connected if they are within $R_0 = 10.5$Å; $l_{ij}$ is the distance between the nodes in the pdb structure; the force constants are $k_{ij} = 189$ Kcal/Mol for sequence neighbors and $k_{ij} = 4.5$ Kcal/Mol otherwise. I placed nodes at the side-chain geometric centers (except for glycine, which has no side chain, for which I used $C_\alpha$ coordinates).

**LFENM perturbations** A mutation is emulated by perturbing independently each of the springs connected to the mutated site by adding perturbations $\delta l_{ij}$. Here, I used $\delta l_{ij} \sim N(0, \sigma = 0.3\text{Å})$.

**Calculation of model parameters $a_S$ and $a_A$**

**LFENM mutational scan** To calculate model rates for a given protein, I started by performing a full mutational scan. For each enzyme, I used its pdb structure to built the ENM. Then, at each site I introduced $N = 19$ LFENM mutations and, for each mutation, I calculated $\Delta\Delta G$ (Eq. S28) and $\Delta\Delta G^*$ (Eq. S45).

**Fitting of parameters $a_S$ and $a_A$** Then, I calculated model rates $K(r)$ using Eq. S13. The model parameters $a_S$ and $a_A$ were obtained by minimizing the Residual Sum of Squares $RSS = \sum_r (K_{\text{obs}}(r) - K(r))^2$ using the general purpose optimization function `optim` of `R` package **stats**.

**Calculation of $< \Delta\Delta G^* >$ and $1 - \omega_A$** Finally, I averaged over mutations at each site to obtain the site-specific physical coupling, $< \Delta\Delta G^* >$, and evolutionary coupling $1 - \omega_A$.

#### 1.3 Data set

I studied data set of 157 enzymes, a subset of the 524 enzymes used in [5]. Specifically, I kept only monomeric enzymes and removed those that had missing amino acids or broken chains, which could result in wrong elastic network models. I also discarded 3 proteins for which the non-linear fitting of the model parameters did not converge. Catalytic residue information was obtained from the Catalytic Site Atlas [6]. The structures of these proteins were obtained from the RCSB protein database[7]. The data set is diverse: no two enzymes have more than 25% sequence identity and there are representatives of the main SCOP structural classes [8] and of the six main EC functional classes [9]. For some details of the data set, see Figure S1

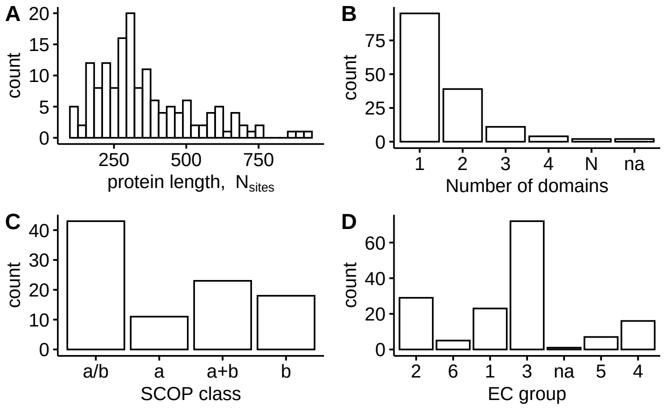


Figure S1: **Properties of the enzymes studied.** The data set is composed of 157 monomeric enzymes. **A**: distribution of sequence lengths. **B**: distribution of number of domains. **C**: distribution of SCOP class for mono-domain cases. **D**: distribution of EC group, the first classification level of the Enzyme Commission number.

#### 1.4 Empirical rates

I used site-specific "observed" rates $K_{\text{obs}}$ calculated before by Jack et al [5]. Briefly, estimating $K_{\text{obs}}$ involves finding homologous sequences, aligning them, inferring a phylogeny, and using the sequence alignment and phylogeny as input for the program `rate4site` to estimate the substitution rate of each site [10].

### 2 Supplementary Results

#### 2.1 MSA assessment

The model MSA predicts site-specific evolutionary rates from mutational changes of free energy ($\Delta\Delta G$), and activation free energy ($\Delta\Delta G^*$). MSA was extensively assessed against other models and empirical data in [1] where it was shown to fit empirical rates and their variation with distance from the active site. For completeness, a brief assessment is replicated here. Fig. S2 shows that the correlation coefficient between predicted and observed rates varies in the range $0.19 <= R <= 0.80$, with an average of $R = 0.62$, which demonstrates the excellent agreement between predicted and empirical rates.

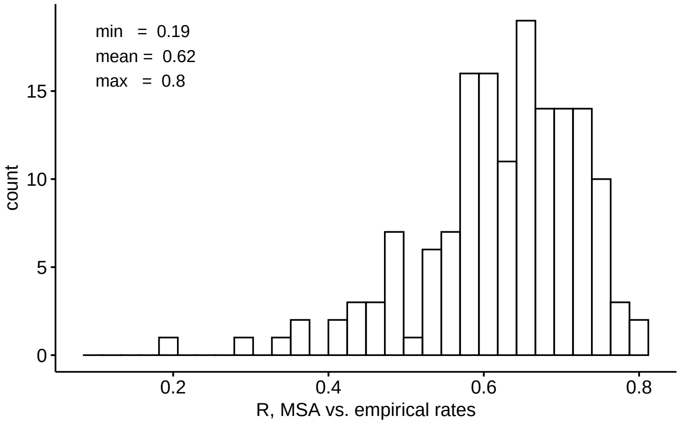


Figure S2: **MSA rates vs. empirical rates.** Distribution of the correlation coefficient $R$ between rates predicted by the model MSA and empirical rates, inferred from multiple sequence alignments using the program Rate4Site by Jack et al. [5].

#### 2.2 On the use of LFENM to calculate $\Delta\Delta G$ and $\Delta\Delta G^*$

In this section, I consider three possible caveats related to using the Linearly Forced Elastic network model (LFENM) to calculate $\Delta\Delta G$ and $\Delta\Delta G^*$. First, *a priori*, a possible limitation of LFENM is that being a coarse grained model, it might not predict $\Delta\Delta G$ and $\Delta\Delta G^*$ accurately. However, In previous studies I have shown that predictions performed using a stability-based model, MS, with LFENM $\Delta\Delta G$ gives very similar results to the presumably better all-atom method FoldX, that has been especially parametrised to calculate $\Delta\Delta G$ values. This is confirmed by Fig. S3, that shows that there is no significant difference between the fit to empirical of rates obtained with LFENM or FoldX. If anything LFENM gives slightly better results.

Second, a related issue is whether LFENM is able to predict $\Delta\Delta G^*$. At this point, I am unable to fully demonstrate it can. However, the MSA model, which uses LFENM-calculated $\Delta\Delta G$ and $\Delta\Delta G^*$, clearly outperforms stability-based models that use either LFENM $\Delta\Delta G$ of FoldX $\Delta\Delta G$ (Fig. S3). In addition, stability-only models *fail* to reproduce the observed dependence of evolutionary rate with increasing distance from the active site, while the MSA model, by including LFENM $\Delta\Delta G^*$, succeeds in predicting the rate-distance dependence quantitatively [1]. Therefore, LFENM $\Delta\Delta G^*$ values seem to capture how coupling varies with distance, which is the interest of the present paper.

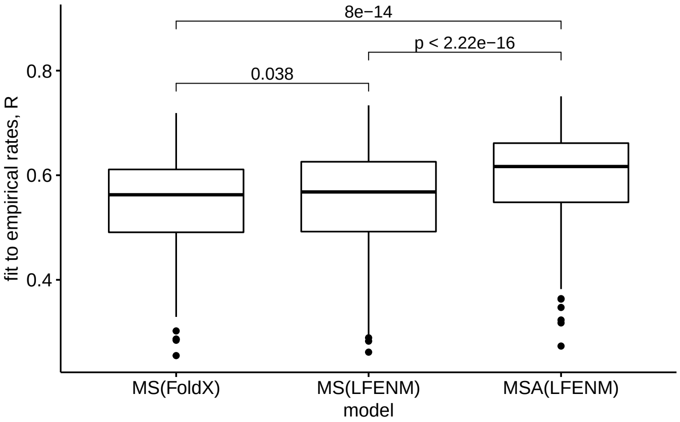


Figure S3: **Comparison of three models: MSA(LFENM), MS(LFENM), and MS(FoldX).** MSA(LFENM) is model MSA that uses $\Delta\Delta G$ and $\Delta\Delta G^*$ calculated using the LFENM coarse-grained model; MS(LFENM) is a stability-only model that uses LFENM $\Delta\Delta G$ values; MS(FoldX) is the same stability model but based on FoldX $\Delta\Delta G$ values. R is the correlation coefficient between model predictions and empirical rates. Means are compared Wilcoxon's signed-rank test for paired cases.

A final limitation to discuss is that LFENM models random perturbations, not actual amino-acid mutations. In elastic network models, sequence is not explicit, but implicit in the protein's conformation. A wild-type node state $\gamma_0$ is implicit in $\mathbf{r}^0_{\text{wt}}$, and a mutant state $\gamma$ is implicit in $\mathbf{r}^0_{\text{mut}}$. Since mutants are obtained using random perturbations, these node states cannot be mapped to actual natural amino acids. Therefore, $\Delta\Delta G$ and $\Delta\Delta G^*$ calculated using LFENM do not correspond to actual amino-acid mutations. All the same, LFENM $\Delta\Delta G$ and $\Delta\Delta G^*$ do get the statistics correctly. MSA rates, which are averages over perturbations, are in very good agreement with observed rates, which are averages over actual amino-acid mutations [1]. Thus, LFENM is suitable for the study of the variation of rates among sites.

#### 2.3 On the meaning of parameter $a_A$ and its huge variation

As shown in Figure 2 of the main text, evolutionary coupling range varies widely among enzymes, from 1.9 Å to 19.7 Å, as a result of a huge variation of parameter $a_A$ over 4 orders of magnitude, from $6 \times 10^{-2}$ to $2 \times 10^3$. Why does $a_A$ vary so widely? The MSA model assumes that the only relevant enzyme property is $\Delta\Delta G^*$, and that $a_A$ represents external selection pressure. However, there might be some enzyme property missed by $\Delta\Delta G^*$ that influences $a_A$. In other words, $a_A$ could be affected by internal or external factors. As possible internal factors, I considered several properties related to protein size, shape, or architecture. I found that $a_A$ is independent of protein length, average solvent accessibility, average packing density, number of domains, and SCOP structural class (see Figure S4). This independence from internal properties suggests that the huge variation of $a_A$ among enzymes probably reflects a variation of *external* factors affecting enzyme evolution; $a_A$ would represent external selection pressure on catalytic activity. Therefore, I take $a_A$ to stand for external functional selection pressure. It's large variation among enzymes would be consistent with the finding that functional constraints related to enzyme metabolic roles vary very widely [11].

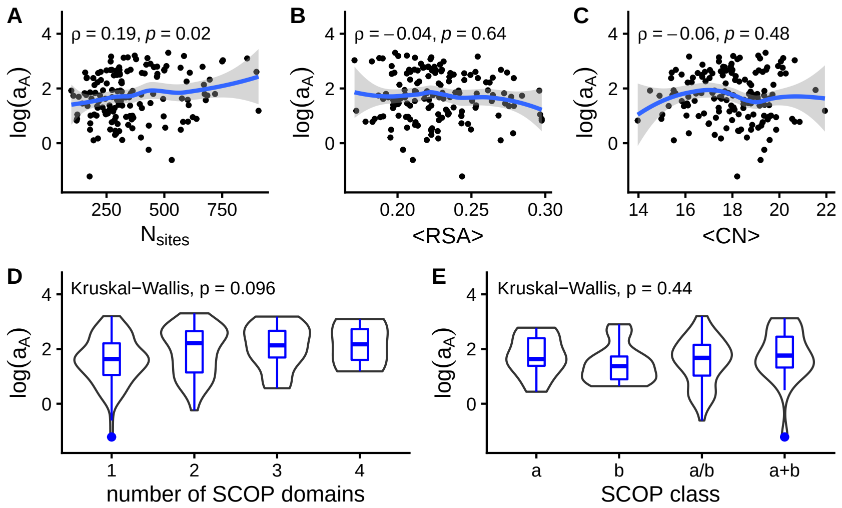


Figure S4: **Non-dependence of $a_A$ on internal enzyme properties.** Model parameter $a_A$ vs. various internal properties: **A**: protein length, $N_{\text{nsites}}$; **B**: average Relative Solvent Accessibility ($< RSA >$), which increases with protein size, depending on shape; **C**: average Contact Number ($< CN >$), a property of the contact network that decreases with size; **D**: number of SCOP domains; **E**: SCOP structural class. $\rho$ is Spearman's correlation coefficient; $p$ stands for p-value; the Kruskal-Wallis test was used to compare means.

#### 2.4 Evolutionary coupling as non-linearly amplified physical coupling

As described in the main text, I found that while physical coupling displays a very similar short-range exponential decrease with distance from the active residues, the distance-dependence of evolutionary coupling varies widely among enzymes, depending on functional selection pressure. This is due to the non-linear dependence of evolutionary coupling on physical coupling (see Eq. S20 and Figure S5). For very small $a_A$, Eq. S20 can be expanded to first order, which leads to: $1-\omega_A \approx a_A < \Delta\Delta G^* >$, so that at very low selection pressures both couplings are short-range exponential declines. For larger $a_A$, the short-range exponential physical coupling is non-linearly mapped into a long-range sigmoidal evolutionary coupling. Therefore, long-range evolutionary coupling is due to the non-linear amplification of physical coupling under strong functional selection pressure.

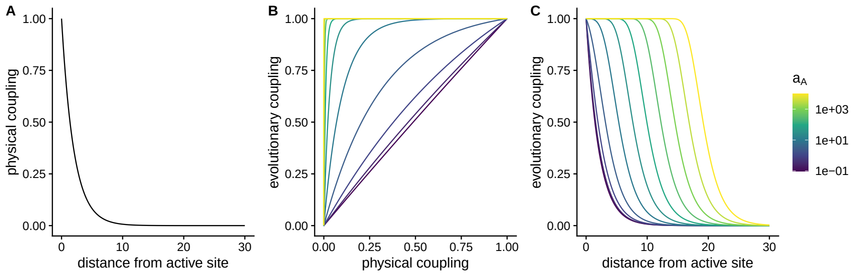


Figure S5: **Selection pressure controls how physical coupling is mapped into evolutionary coupling.** **A**: typical exponentially decreasing physical coupling. **B**: evolutionary coupling vs. physical coupling for increasing functional selection pressure (Eq. S20: $1-\omega_A = 1- < \min(1, e^{-a_A \Delta\Delta G^*}) >$). **C**: distance-dependence of evolutionary coupling, which is the composition of panels **A** and **B**. Note that the same short-range physical coupling (panel **A**) is mapped into evolutionary couplings with increasingly longer range as functional selection pressure increases (panel **C**). The curves of panels **B** and **C** are coloured according the functional selection pressure parameter $a_A$.

### 3 Theory

This section describes the Stability-Activity Model of enzyme evolution, MSA. The present material is adapted from the article where the model was presented, where the reader is referred to for a more detailed description, assessment and discussion [1]. The section is structured as follows. First, I derive the model. Specifically, I pose the fitness landscape (Figure S6A); derive the fixation probability (Figure S6B); derive a mean-field approximation to this fixation probability (Figure S6C); derive the formulas needed to calculate site-specific amino-acid substitution rates (Eq. S13); and derive the function that relates evolutionary coupling to physical coupling (Eq. S20.) Second, I derive a formula to calculate stability changes $\Delta\Delta G$ using the "Linearly-Forced Elastic Network Model" (LFENM) (Eq. S28 and Figure S6D). Finally, I derive a formula to calculate mutational activation free energy changes $\Delta\Delta G^*$ using LFENM (Eq. S45 and Figure S6D).

#### 3.1 The MSA model

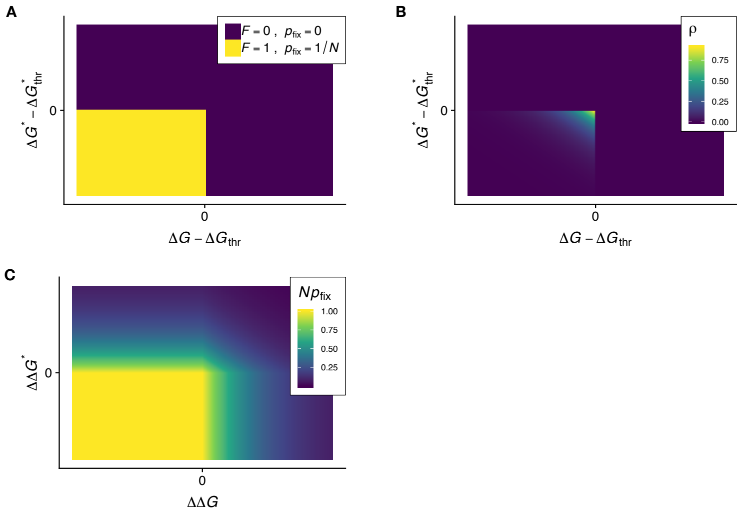


Figure S6: **The stability-activity model MSA.** **A**: Fitness is a 2D step function: enzymes are viable ($F = 1$) and may become fixed ($p^{\text{fix}} = 1/N$, $N$ being population size) only if their folding free energy $\Delta G$ is more negative than a threshold energy $\Delta G_{\text{thr}}$ and if their activation energy barrier $\Delta G^*$ is lower than a threshold $\Delta G^*_{\text{thr}}$. **B**: Distribution of $\Delta G$ and $\Delta G^*$ at evolutionary equilibrium. **C**: A mean field $p^{\text{fix}}$ that depends only on $\Delta\Delta G$ (mutational change of stability) and $\Delta\Delta G^*$ (mutational change of activation energy) is obtained by integrating the $p^{\text{fix}}$ of panel A over the distribution of panel B, . This figure is part of a figure used before in [1].

**Fitness landscape**

The stability-activity model of enzyme evolution MSA is specified by a fitness function that depends on enzyme stability, quantified by the folding free energy $\Delta G$, and activity, quantified by the activation energy of the enzymatic reaction $\Delta G^*$. Specifically, fitness is given by the following two-dimensional step function (Figure S6A):

$$\boxed{F(\Delta G, \Delta G^*) = \begin{cases} 1 & \text{if } \Delta G < \Delta G_{\text{thr}} \text{ and } \Delta G^* < \Delta G^*_{\text{thr}} \\ 0 & \text{if } \Delta G >= \Delta G_{\text{thr}} \text{ or } \Delta G^* >= \Delta G^*_{\text{thr}} \end{cases}} \quad \text{(S1)}$$

Thus, all genotypes with stability above a stability threshold ($\Delta G < \Delta G_{\text{thr}}$) and activation energy barrier below an activation energy threshold ($\Delta G^* < \Delta G^*_{\text{thr}}$) are equally viable ($F = 1$) and genotypes that are too unstable ($\Delta G^* >= \Delta G^*_{\text{thr}}$) and/or too inactive ($\Delta G^* >= \Delta G^*_{\text{thr}}$) are inviable ($F = 0$).

**Fixation probability**

Assuming that the mutation rate $\mu$ is so small that the time between mutations is much larger than the time needed to fix or discard a new mutation, evolution is an origination-fixation process in which a new mutant is either lost or fixed before a another mutant originates [12]. Consider a monoclonal population of $N$ identical haploid genotypes with stability $\Delta G$ and activation energy $\Delta G^*$. For the fitness function of Eq. S1, when a mutant originates, the probability that it becomes fixed, replacing all the ancestor genotypes, is given by:

$$\boxed{p^{\text{fix}} = \begin{cases} \frac{1}{N} & \text{for } \Delta G + \Delta\Delta G < \Delta G_{\text{thr}} \text{ and } \Delta G^* + \Delta\Delta G^* < \Delta G^*_{\text{thr}} \\ 0 & \text{if } \Delta G + \Delta\Delta G >= \Delta G_{\text{thr}} \text{ or } \Delta G^* + \Delta\Delta G^* >= \Delta G^*_{\text{thr}} \end{cases}} \quad \text{(S2)}$$

where $\Delta G + \Delta\Delta G = \Delta G_{\text{mut}}$ is the mutant's stability and $\Delta G^* + \Delta\Delta G^* = \Delta G^*_{\text{mut}}$ is its activation free energy. If the mutant is within the allowed stability and activity thresholds, the mutation is neutral, the mutant has the same fitness as the other $N - 1$ individuals of the population, thus it will become fixed with probability $p^{\text{fix}} = 1/N$. On the other hand, if the mutant is not stable or active enough, it will necessarily become lost, thus $p^{\text{fix}} = 0$.

**Mean-field fixation probability**

It is usually easier to calculate relative free energy changes, such as $\Delta\Delta G$ and $\Delta\Delta G^*$, than absolute values such as $\Delta G$ and $\Delta G^*$. Therefore, for tractability, I follow [13, 14] and derive a *mean field* approximation to the fixation probability that depends only on $\Delta\Delta G$ and $\Delta\Delta G^*$. Specifically, I assume that $\Delta\Delta G$ and $\Delta\Delta G^*$ are independent of the specific sequence background in which the mutation happens. Then, I average the fixation probability over $\Delta G$ and $\Delta G^*$:

$$p^{\text{fix}}(\Delta\Delta G, \Delta\Delta G^*) = \int_{-\infty}^{\infty}\int_{-\infty}^{\infty} p^{\text{fix}}(\Delta G_{\text{wt}}, \Delta G^*_{\text{wt}}, \Delta\Delta G, \Delta\Delta G^*)\rho_S(\Delta G_{\text{wt}})\rho_A(\Delta G^*_{\text{wt}})d\Delta G_{\text{wt}}d\Delta G^*_{\text{wt}} \tag{S3}$$

where $\rho_S$ is the stationary distribution of stabilities and $\rho_A$ the distribution of activation energies. These distributions may be approximated using truncated exponentials (Figure S6B). For evolution on a stability-dependent step-function fitness function, in the rare mutation limit ($\mu N << 1$), the stationary stability distribution is well approximated by a truncated exponential[13]:

$$\rho_S(\Delta G) = \begin{cases} a_S e^{a_S(\Delta G - \Delta G_{\text{thr}})} & if \Delta G < \Delta G_{\text{thr}} \\ 0 & if \Delta G >= \Delta G_{\text{thr}} \end{cases} \tag{S4}$$

Since evolution in the $\Delta G^*$ dimension is mathematically equivalent, I further assume:

$$\rho_A(\Delta G^*) = \begin{cases} a_A e^{a_A(\Delta G^* - \Delta G^*_{\text{thr}})} & if \Delta G^* < \Delta G^*_{\text{thr}} \\ 0 & if \Delta G^* >= \Delta G^*_{\text{thr}} \end{cases} \tag{S5}$$

Replacing Eq. S2, Eq. S4, and Eq. S5 into Eq. S3 and integrating, it follows that

$$\boxed{p^{\text{fix}}(\Delta\Delta G, \Delta\Delta G^*) = 1/N \min(1, e^{-a_S \Delta\Delta G}) \min(1, e^{-a_A \Delta\Delta G^*})} \tag{S6}$$

This is mean field approximation to the fixation probability in terms of mutational changes of stability, $\Delta\Delta G$, and activation energy, $\Delta\Delta G^*$ (see Figure S6C).

##### Substitution rates

The variation of evolutionary constraints within proteins is commonly quantified using site-specific substitution rates. In this section, I derive the formulas needed to calculate site-specific amino-acid substitution rates. First, let us derive site-specific substitution rates as a function of stability and activity changes. The the rate of $\alpha \to \beta$ substitutions (the number of substitutions per unit time) at site $r$ is given by [15]:

$$Q_{\beta\alpha}(r) = N\mu p^{\text{fix}}_{\beta\alpha}(r) \tag{S7}$$

where $N$ is the population size, $\mu$ is the mutation rate, and $p^{\text{fix}}_{\beta\alpha}(r)$ is the probability of fixating the $\alpha \to \beta$ mutation at site $r$, given by Eq. S6:

$$p^{\text{fix}}_{\beta\alpha}(r) = \frac{1}{N} \min(1, e^{-a_S \Delta\Delta G_{\beta\alpha}(r)}) \min(1, e^{-a_A \Delta\Delta G^*_{\beta\alpha}(r)}) \tag{S8}$$

where $\Delta\Delta G_{\beta\alpha}(r)$ is the free energy change due to the $\alpha \to \beta$ mutation at site $r$, and $\Delta\Delta G^*_{\beta\alpha}(r)$ the corresponding activation energy change. Replacing Eq. S8 into Eq. S7, it follows:

$$Q_{\beta\alpha}(r) = \mu \min(1, e^{-a_S \Delta\Delta G_{\beta\alpha}(r)}) \min(1, e^{-a_A \Delta\Delta G^*_{\beta\alpha}(r)}) \tag{S9}$$

Second, I derive the site-specific amino-acid probability distributions at evolutionary equilibrium. Let $\pi_\gamma(r)$ be the probability of finding amino acid $\gamma$ at site $r$ at a certain evolutionary time $t$. These probabilities satisfy the evolution equations:

$$\frac{d\pi_\beta(r)}{dt} = \sum_\alpha Q_{\beta\alpha}(r)\pi_\alpha(r). \tag{S10}$$

At equilibrium, ${d\pi_\gamma(r)}/{dt} = 0$. From this condition and Eq. S9, it can be shown that the equilibrium probabilities are given by:

$$\pi_\alpha^{eq}(r) = \frac{e^{-a_S \Delta\Delta G_{\alpha\gamma_0}(r) - a_A \Delta\Delta G^*_{\alpha\gamma_0}(r)}}{\sum_\gamma e^{-a_S \Delta\Delta G_{\gamma\gamma_0}(r) - a_A \Delta\Delta G^*_{\gamma\gamma_0}(r)}} \tag{S11}$$

where $\pi_\alpha^{eq}(r)$ is the probability of finding amino acid $\alpha$ at site $r$ at evolutionary equilibrium, $\gamma_0$ stands for a reference state (amino acid) that may be chosen arbitrarily without loss of generality, and the denominator ensures that $\sum_\alpha \pi_\alpha^{eq}(r) = 1$.

Finally, I derive the total site-specific substitution rates. At evolutionary equilibrium, the total substitution rate of site $r$ is obtained by summing over final states and averaging over initial states, which gives:

$$K(r) = \sum_\alpha \sum_{\beta \neq \alpha} Q_{\beta\alpha}(r)\pi_\alpha^{eq}(r) \tag{S12}$$

where $Q_{\beta\alpha}(r)$ is given by Eq. S9 and $\pi_\alpha^{eq}(r)$ is given by Eq. S11. Therefore, replacing Eq. S9 and Eq. S11 into Eq. S12, it follows that the rate of substitutions at a site $r$ is given by:

$$K(r) = \sum_\alpha \sum_{\beta \neq \alpha} Q_{\beta\alpha}(r)\pi_\alpha^{eq}(r) \tag{S13a}$$

$$Q_{\beta\alpha}(r) = \mu \min(1, e^{-a_S \Delta\Delta G_{\beta\alpha}(r)}) \min(1, e^{-a_A \Delta\Delta G^*_{\beta\alpha}(r)}) \tag{S13b}$$

$$\pi_\alpha^{eq}(r) = \frac{e^{-a_S \Delta\Delta G_{\alpha\gamma_0}(r) - a_A \Delta\Delta G^*_{\alpha\gamma_0}(r)}}{\sum_\gamma e^{-a_S \Delta\Delta G_{\gamma\gamma_0}(r) - a_A \Delta\Delta G^*_{\gamma\gamma_0}(r)}} \tag{S13c}$$

Using this equation, given selection parameters $a_S$ and $a_A$, site-specific rates may be calculated from mutational changes of stability, $\Delta\Delta G_{\beta\alpha}$, and activation free energy, $\Delta\Delta G^*_{\beta\alpha}$.

##### Relative rate

Let $K$ be the rate of evolution of a protein or a protein site and $K_0$ be the evolutionary rate in the neutral evolution case (i.e. mutations are neutral, or, equivalently, when natural selection does not operate). Then, the relative rate of evolution is defined by:

$$\omega \equiv \frac{K}{K_0} \tag{S14}$$

The value of $\omega$ is related natural selection. $\omega > 1$ for positive selection (adaptive evolution), and $\omega < 1$ for negative/purifying selection.

For the MSA model, the relative slowdown is obtained by replacing Eq. S13 into definition Eq. S14 (note that $K_0 = K(a_S = 0, a_A = 0)$), which leads to:

$$\omega(r) = \frac{1}{N_{aa} - 1} \sum_{\alpha} \sum_{\beta \neq \alpha} \min(1, e^{-a_S \Delta\Delta G_{\beta\alpha}(r)}) \min(1, e^{-a_A \Delta\Delta G^*_{\beta\alpha}(r)}) \pi_\alpha(r)^{eq} \tag{S15}$$

where $\pi_\alpha(r)$ is the probability of finding amino-acid $\alpha$ in residue $r$ at equilibrium and $N_{aa}$ is the number of possible amino acids (commonly 20).

The notation can be simplified by noting that for any property $X$ that depends on mutations $\alpha \to \beta$, the average over an evolutionary trajectory is given by:

$$\langle X \rangle = \frac{1}{N_{aa} - 1} \sum_{\alpha} \sum_{\beta \neq \alpha} X_{\beta\alpha} \pi_\alpha^{eq} \tag{S16}$$

Therefore, using this notation in Eq. S15, gives the more compact:

$$\omega(r) = \langle \min(1, e^{-a_S \Delta\Delta G(r)}) \min(1, e^{-a_A \Delta\Delta G^*(r)}) \rangle \tag{S17}$$

This equation relates mutational changes $\Delta\Delta G$ and $\Delta\Delta G^*$ to relative rates of evolution $\omega$ for the model MSA. That MSA is a model of evolution under negative selection can be seen in Figure S7. $\omega$ is maximum at $a_S = a_A = 0$, when selection does not operate, and it decreases when the selection-pressure parameters $a_S$ and/or $a_A$ increase.

**Relative slowdown**

To measure the effects of selection pressure, rather than $\omega$, it is more convenient to use $1 - \omega$, which is the *relative slowdown*:

$$1 - \omega = \frac{K_0 - K}{K_0} \tag{S18}$$

For instance, a protein, or a protein site, with $\omega = 0.7$ is evolving at 70% of its unselected rate; its slowdown is $1 - \omega = 0.3$, which means that it is evolving 30% more slowly than it would be under no selection. Replacing Eq. S17 into Eq. S18, it follows that:

$$\boxed{1 - \omega(r) = 1 - \langle \min(1, e^{-a_S \Delta\Delta G(r)}) \min(1, e^{-a_A \Delta\Delta G^*(r)}) \rangle} \tag{S19}$$

This expresses the slowdown of the evolution of a protein site as a function of the mutational changes of stability ($\Delta\Delta G$) and activation free energy ($\Delta\Delta G^*$), depending on parameters $a_S$, which measures selection pressure on stability, and $a_A$, which measures selection pressure on activity.

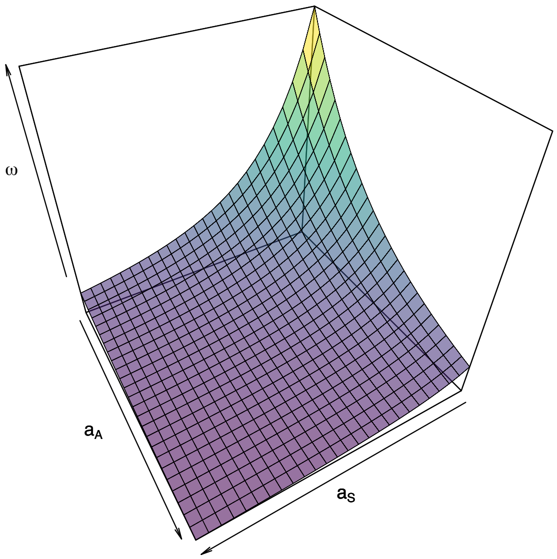


Figure S7: **Variation of the relative rate $\omega$ with selection pressure.** Plot of the relative rate $\omega$ predicted by MSA for one protein site of the protein with pdb code 1PMI, as a function of the model's selection parameters $a_S$ and $a_A$. When $a_S = a_A = 0$, there is no selection and the rate is maximum. As selection on stability ($a_S$) or on activity ($a_A$) increase, $\omega$ decreases.

##### Slowdown due selection on activity

Eq. S19 relates the slowdown of the evolution of a site to the effects of mutating this site on stability and activity. To study the effects of selection on activity, I consider a hypothetical scenario in which selection on stability is turned off. Replacing $a_S = 0$ into Eq. S19 it follows that:

$$\boxed{1 - \omega_A(r) = 1 - \langle \min(1, e^{-a_A \Delta\Delta G^*(r)}) \rangle} \tag{S20}$$

where $1 - \omega_A(r)$ is the slowdown of site $r$ due to selection on activity. Note that $\omega_A = \omega(a_S = 0, a_A)$, so that it the curve at the intersection of the $omega(a_S, a_A)$ surface and the $a_A - \omega$ plane (see Figure S7).

##### Physical and evolutionary couplings

I will show below that $\Delta\Delta G^*(r)$ results from modifications at the active-site location due to mutations at residue $r$ [1]. Therefore, $\Delta\Delta G^*(r)$ represents the *physical coupling* between residue $r$ and the enzyme's active site. Accordingly, this physical coupling causes a slowdown $1 - \omega_A(r)$, which, therefore, represents the *evolutionary coupling* between residue $r$ and the active site. Thus,

Eq. S20 governs how evolutionary coupling $(1 - \omega_A)$ depends on physical coupling $(\Delta\Delta G^*)$ and functional selection pressure $(a_A)$.

#### 3.2 LFENM calculation of $\Delta\Delta G$

The site-specific rates of evolution that MSA aims to predict depend on mutational changes of stability, $\Delta\Delta G$, and of activation energy, $\Delta\Delta G^*$. These are represented in Figure S8. To calculate these mutational changes, I use the Linearly Forced Elastic Network Model (LFENM) [2, 3, 16, 17]. This section describes the LFENM model and its use to calculate $\Delta\Delta G$.

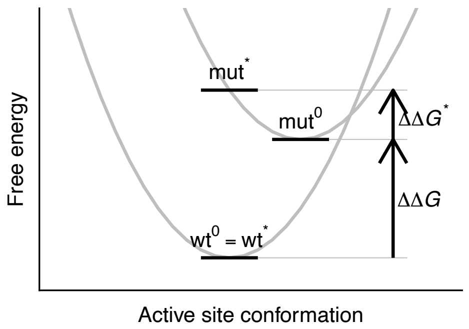


Figure S8: **Mutational impact on stability and activity.** $\Delta\Delta G$ is the difference between energy minima; $\Delta\Delta G^*$ is the energy needed to distort the mutant from its equilibrium conformation $\mathrm{mut}^0$ to the active conformation $\mathrm{mut}^*$ (the wild type's active site is assumed preorganized in the active conformation: $\mathrm{wt}^0 = \mathrm{wt}^*$).

##### Wild type elastic network

The LFENM model represents proteins as networks of nodes, representing amino acids, connected by harmonic springs, representing interactions. The energy function of a given wild type protein is given by

$$V_{\mathrm{wt}} = \frac{1}{2}\sum_{i<j} k_{ij}(d_{ij} - l_{ij})^2 \tag{S21}$$

where $d_{ij}$ is the distance between nodes $i$ and $j$ in an arbitrary conformation, and $k_{ij}$ and $l_{ij}$ are, respectively, the force constant and equilibrium length of the spring connecting nodes $i$ and $j$. $V_{\mathrm{wt}}$ is usually approximated using a second-order Taylor expansion of Eq. S21:

$$V_{\mathrm{wt}}(\mathbf{r}) = \frac{1}{2}(\mathbf{r} - \mathbf{r}^0_{\mathrm{wt}})^T\mathbf{K}(\mathbf{r} - \mathbf{r}^0_{\mathrm{wt}}) \tag{S22}$$

where $T$ stands for 'transpose', $\mathbf{r}$ is the column vector of node coordinates that represents and arbitrary conformation, $\mathbf{r}^0_{\text{wt}}$ is the wild type's equilibrium conformation, and $\mathbf{K}$ is the Hessian matrix of second derivatives of $V_{\text{wt}}$.

##### Mutated elastic network

The LFENM model simulates mutations by perturbing the equilibrium length of the springs that connect the mutated site to its neighbors. Thus, introducing a mutation at site $r$ results in a mutant with energy

$$V_{\text{mut}} = \frac{1}{2}\sum_{i<j} k_{ij}(d_{ij} - l_{ij} - \delta l_{ij})^2 \tag{S23}$$

where $\delta l_{ij}$ is a random perturbation added to $l_{ij}$.(Only contacts of site $r$ are perturbed: $\delta l_{ij} \neq 0$ only for $i = r$ or $j = r$, and $\delta l_{ij} = 0$ otherwise.)

In general, the mutation will shift the equilibrium conformation. The mutant's equilibrium structure is given by [2]:

$$\mathbf{r}^0_{\text{mut}} = \mathbf{r}^0_{\text{wt}} + \delta\mathbf{r}^0, \tag{S24}$$

where the mutational change of the equilibrium structure is given by

$$\delta\mathbf{r}^0 = \mathbf{r}^0_{\text{mut}} - \mathbf{r}^0_{\text{wt}} = -\mathbf{K}^{-1}\mathbf{f}, \tag{S25}$$

where $\mathbf{f}$ is a "force´´ vector that depends on the perturbations $\delta l_{ij}$. (perturbing the spring-length parameter $l_{ij}$ by adding $\delta l_{ij}$ is equivalent to applying a force of magnitude $f_{ij}$ in the direction of the vector that goes from site $i$ to site $j$.)

Expanding the mutant's energy up to second order around its equilibrium conformation $\mathbf{r}^0_{\text{mut}}$ leads to

$$V_{\text{mut}}(\mathbf{r}) = V^0_{\text{mut}} + \frac{1}{2}(\mathbf{r} - \mathbf{r}^0_{\text{mut}})^T\mathbf{K}(\mathbf{r} - \mathbf{r}^0_{\text{mut}}) \tag{S26}$$

where

$$V^0_{\text{mut}} = \frac{1}{2}\sum_{j\sim i} k_{ij}\delta l^2_{ij} - \frac{1}{2}(\delta\mathbf{r}^0)^T\mathbf{K}\delta\mathbf{r}^0 \tag{S27}$$

is the mutant's minimum energy.

Note that $\mathbf{K}$ in the previous equations is the Hessian matrix of $V_{\text{wt}}$ and $V_{\text{mut}}$; since $\mathbf{K}$ is not affected by *linear* perturbations, $\mathbf{K}_{\text{mut}} = \mathbf{K}_{\text{wt}} = \mathbf{K}$. For more details on the LFENM see [2, 3].

##### LFENM formula for $\Delta\Delta G$

Consider the free energy change due introducing a single mutation in the wild-type protein. Let $\gamma_0$ be the state of node $r$ of the elastic network that represents the wild-type protein; and let $\gamma$ represent the mutated state of node $r$ of the elastic network representing the mutant protein, obtained through perturbing the network, as explained above. The free-energy change due to the mutation,

$\Delta\Delta G_{\gamma\gamma_0}(r)$, is the free energy difference between the mutated elastic network and the wild type elastic network (Figure S8). Using basic statistical physics, from Eq. S22, Eq. S26, and Eq. S27, it can be found that:

$$\boxed{\Delta\Delta G_{\gamma\gamma_0}(r) = \frac{1}{2}\sum_{j<i} k_{ij}\delta l_{ij}^2 - \frac{1}{2}(\mathbf{r}^0_{\text{mut}} - \mathbf{r}^0_{\text{wt}})^T \mathbf{K}(\mathbf{r}^0_{\text{mut}} - \mathbf{r}^0_{\text{wt}})} \tag{S28}$$

where $\mathbf{r}^0_{\text{mut}} - \mathbf{r}^0_{\text{wt}}$ may be obtained using Eq. S25 and $\delta l_{ij}$ are the perturbations added to the springs of the mutated site $r$ to model the mutation $\gamma_0 \to \gamma$. The free-energy change for an arbitrary mutation $\alpha \to \beta$ is given by:

$$\Delta\Delta G_{\beta\alpha}(r) = \Delta\Delta G_{\beta\gamma_0}(r) - \Delta\Delta G_{\alpha\gamma_0}(r) \tag{S29}$$

This follows from basic Thermodynamics and the fact that a mutation $\alpha \to \beta$ can be obtained via the path $\alpha \to \gamma_0 \to \beta$.

N.B: mutations of the wild-type protein are always destabilising ($\Delta\Delta G_{\gamma\gamma_0}(r) >= 0$.) By construction, all springs of the wild-type elastic network are relaxed. As a result of the mutation, the springs connected to the mutated site become stressed, so that in the wild-type conformation $\mathbf{r}^0_{\text{wt}}$ these springs are either too compressed or too extended. This local stress is represented by the first term of of Eq. S28, which is always positive. Part of this stress is relaxed in the mutant conformation $\mathbf{r}^0_{\text{mut}}$, which is represented by the negative second term of of Eq. S28. However, while the local stress near the mutation is now somewhat reduced, other springs move out of their ideal equilibrium lengths, so that even in the mutant's equilibrium conformation all springs are stressed. Therefore, $\Delta\Delta G_{\gamma,\gamma_0}(r) >= 0$ for all $r$ and $\gamma$. In contrast, an arbitrary $\alpha \to \beta$ mutation may be destabilizing or stabilizing depending on the relative magnitudes of the two terms of Eq. S29 [1].

#### 3.3 LFENM calculation of $\Delta\Delta G^*$

**Activation energy barrier $\Delta G^*$**

Consider the following simple mechanism of enzyme catalysis:

$$\text{E} + \text{S} \rightleftharpoons \text{ES} \rightarrow ES^* \rightarrow \text{E} + \text{P} \tag{S30}$$

The first reaction is the binding between enzyme and substrate to form the complex $ES$. The second reaction is the formation of products via a transition state $ES^*$. According to transition state theory, for low substrate concentrations this reaction has second order kinetics with rate constant [18]

$$k = \frac{k_B T}{h} e^{-\frac{1}{k_B T}\Delta G^*} \tag{S31}$$

where $k_B$ is Boltzmann's constant, $T$ is the absolute temperature, $h$ is Planck's constant, and

$$\Delta G^* = G(ES^*) - G(E) - G(S) \tag{S32}$$

is the *activation free energy*, which is the free-energy difference between the enzyme-substrate activated complex and enzyme and substrate in their free solution states.

Following [19, 18], we can break the activation process into two steps, a *distortion step*, in which enzyme and substrate adopt the conformation they have in the activated complex, followed by a *vertical binding step*, in which enzyme and substrate in the right *activated poses* bind to each other without further conformational change. Then:

$$\Delta G^* = \Delta G(E \to E^*) + \Delta G(S \to S^*) + \Delta G(E^* + S^* \to ES^*) \tag{S33}$$

The first term is the distortion energy needed for the enzyme to adopt its activated-complex pose, the second is the substrate distortion energy, and the third term is the vertical binding energy, the free energy released when enzyme and substrate in the right poses bind together.

##### $\Delta\Delta G^*$ is the mutant's distortion free energy

This section introduces two reasonable assumptions: (1) that mutations change only the enzyme distortion contribution to $\Delta\Delta G^*$, and (2) that the wild-type's active site is pre-organized into the activated pose. With this assumptions, it follows that $\Delta\Delta G^*$ is the mutant's distortion free energy. (Figure S6D.)

**Assumption: mutations change only the enzyme distortion contribution.** From Eq. S33, the free-energy change due to a mutation can be partitioned as:

$$\Delta\Delta G^* = \Delta\Delta G(E \to E^*) + \Delta\Delta G(S \to S^*) + \Delta\Delta G(E^* + S^* \to ES^*) \tag{S34}$$

The mutation will not affect the distortion energy of the substrate in solution, therefore the second term is zero. The third term should be affected only by mutations of the residues directly involved in binding (i.e. active-site residues and subrate-binding residues). Since here I am interested in exploring the effect of mutating sites distant from the active residues, to explore long-range coupling, the third term will also likely be zero. Therefore, in this work, I assume that mutations affect only the distortion energy of the enzyme, so that only the first term of Eq. S34 contributes to $\Delta\Delta G^*$:

$$\Delta\Delta G^* = \Delta\Delta G(E \to E^*) \ = \Delta G_{\mathrm{mut}}(E \to E^*) - \Delta G_{\mathrm{wt}}(E \to E^*) \tag{S35}$$

**Assumption: the wild-type's active site is perfectly pre-organised.** In general, enzyme active sites are pre-organized: their native conformation is very similar to that observed in the activated complex. Therefore, I further assume that the wild-type's active site is perfectly pre-organized, so that

$$\Delta G_{\mathrm{wt}}(E \to E^*) = 0 \tag{S36}$$

**Result: $\Delta\Delta G^*$ is the mutant's distortion free energy.** Therefore, replacing Eq. S36 into Eq. S35, it follows:

$$\boxed{\Delta\Delta G^* = \Delta G_{\text{mut}}(E \to E^*)} \tag{S37}$$

Thus, the mutational activation energy change is identical to the free energy needed to distort the mutant from its equilibrium conformation to the activated pose.

##### $\Delta\Delta G^*$ from conformational distributions

$\Delta\Delta G^*$ can be obtained from the mutant's conformational distribution. Let $\mathbf{r}_a$ be the column vector of coordinates of the catalytic residues, and $\mathbf{r}_n$ be the vector of coordinates of the rest of the residues. The enzyme distortion free energy is the energy necessary to make the active residues adopt a specific active pose $\mathbf{r}_a^*$, regardless of the conformation of the rest of the sites $\mathbf{r}_n$. Further, since we have assumed that the wild-type protein has a perfectly pre-organised active site, the active conformation is the wild-type's conformation: $\mathbf{r}_a^* = \mathbf{r}_{a,\text{wt}}^0$. Then, from basic statistical physics it follows that the mutant's distortion free energy can be obtained from

$$\Delta G_{\text{mut}}(E \to E^*) = -k_B T \log \frac{\rho_{\text{mut}}(\mathbf{r}_{a,\text{wt}}^0)}{\rho_{\text{mut}}(\mathbf{r}_{a,\text{mut}}^0)} \tag{S38}$$

where $\rho_{\text{mut}}(\mathbf{r}_a)$ is the marginal distribution of $\mathbf{r}_a$ conformations, $\mathbf{r}_{a,\text{wt}}^0)$ is wild-type native conformation of the active residues (which, due to pre-organisation, is the enzyme's active pose,) and $\mathbf{r}_{a,\text{mut}}^0$ is the active site mutant's equilibrium conformation. Replacing Eq. S38 into Eq. S37, gives:

$$\Delta\Delta G^* = -k_B T \log \frac{\rho_{\text{mut}}(\mathbf{r}_{a,\text{wt}}^0)}{\rho_{\text{mut}}(\mathbf{r}_{a,\text{mut}}^0)} \tag{S39}$$

This equation allows the calculation of $\Delta\Delta G^*$ given $\rho_{\text{mut}}(\mathbf{r}_a^*)$.

##### LFENM mutant's conformational distribution

From the previous equation, to finish the derivation of $\Delta\Delta G^*$, we need the mutant's marginal distribution function $\rho_{\text{mut}}(\mathbf{r}_a)$. We can obtain this by integration of the LFENM distributions.

The mutant's conformational distribution is given by the Boltzmann density function $\rho_{mut}(\mathbf{r}) = e^{-\beta V_{\text{mut}}(\mathbf{r})}/Z$ where the partition function $Z$ is a normalization constant. From the LFENM's $V_{\text{mut}}$ given by Eq. S26, it follows that:

$$\rho_{\text{mut}}(\mathbf{r}) = \frac{e^{-1/2(\mathbf{r}-\mathbf{r}_{\text{mut}}^0)^T \mathbf{\Sigma}^{-1}(\mathbf{r}-\mathbf{r}_{\text{mut}}^0)}}{\sqrt{2\pi|\mathbf{\Sigma}|}} \tag{S40}$$

where $\mathbf{\Sigma}$ is the variance-covariance matrix, given by:

$$\mathbf{\Sigma} = \frac{1}{k_B T}\mathbf{K}^{-1} \tag{S41}$$

(Note: $\mathbf{K}$ is singular, thus the pseudo-inverse is used.)

The marginal distribution $\rho_{\mathrm{mut}}(\mathbf{r}_a)$ is obtained by integrating Eq. S40 over the coordinates of non-active residues $\mathbf{r}_n$, which gives:

$$\rho_{\mathrm{mut}}(\mathbf{r}_a) = \frac{e^{-1/2(\mathbf{r}_a - \mathbf{r}^0_{a,\mathrm{mut}})^T \mathbf{\Sigma}^{-1}_{aa}(\mathbf{r}_a - \mathbf{r}^0_{a,\mathrm{mut}})}}{\sqrt{|2\pi\mathbf{\Sigma}_{aa}|}} \tag{S42}$$

where $\mathbf{\Sigma}_{aa}$ is the block of $\mathbf{\Sigma}$ that corresponds to the coordinates of the active-site residues. This Marginal distribution may be thought of as the distribution of of a reduced-dimensionality energy function:

$$V_{\mathrm{mut}}(\mathbf{r}_a) = V_{\mathrm{mut}}(\mathbf{r}^0_{\mathrm{mut}}) + \frac{1}{2}(\mathbf{r}_a - \mathbf{r}^0_{a,\mathrm{mut}})^T \mathbf{K}^{\mathrm{eff}}_{aa}(\mathbf{r}_a - \mathbf{r}^0_{a,\mathrm{mut}}) \tag{S43}$$

where

$$\mathbf{K}^{\mathrm{eff}}_{aa} \equiv k_B T \mathbf{\Sigma}^{-1}_{aa} \tag{S44}$$

This matrix may be interpreted as an effective energy matrix for motion of the active residues that takes into account their direct interactions and indirect interactions via their coupling with non-active residues (see e.g. [20]).

##### LFENM formula to calculate $\Delta\Delta G^*$

Finally, using the LFENM mutant's marginal distribution derived above, we obtain $\Delta\Delta G^*$. Replacing Eq. S42 into into Eq. S39 and using Eq. S44, leads to:

$$\boxed{\Delta\Delta G^*_{\gamma\gamma_0}(r) = \frac{1}{2}(\mathbf{r}^0_{a,\mathrm{wt}} - \mathbf{r}^0_{a,\mathrm{mut}})^T \mathbf{K}^{\mathrm{eff}}_{aa}(\mathbf{r}^0_{a,\mathrm{wt}} - \mathbf{r}^0_{a,\mathrm{mut}})} \tag{S45}$$

This is the activation free energy change for introducing a mutation $\gamma_0 \to \gamma$ into site $r$ of the wild-type protein. For arbitrary mutations $\alpha \to \beta$, $\Delta\Delta G^*$ is given by:

$$\Delta\Delta G^*_{\beta\alpha}(r) = \Delta\Delta G^*_{\beta\gamma_0}(r) - \Delta\Delta G^*_{\alpha\gamma_0}(r) \tag{S46}$$

which follows from the fact that mutation $\alpha \to \beta$ can be obtained by the following sequence: $\alpha \to \gamma_0 \to \beta$.

Note that according to Eq. S45, $\Delta\Delta G^*_{\gamma\gamma_0}(r) >= 0$, so that mutations of the wild-type always increase the activation energy barrier. The reason for this is that the wild type enzyme is assumed to be perfectly pre-organised, having zero distortion contribution to the activation energy barrier. Any mutation will shift the mutant's equilibrium structure away from the ideal active pose, increasing the deformation contribution to the free energy barrier. However, for other mutations $\Delta\Delta G^*_{\beta\alpha}(r)$, given by Eq. S46, may be positive or negative.
